# TripletProt: Deep Representation Learning of Proteins based on Siamese Networks

**DOI:** 10.1101/2020.05.11.088237

**Authors:** Esmaeil Nourani, Ehsaneddin Asgari, Alice C. McHardy, Mohammad R.K. Mofrad

## Abstract

We introduce TripletProt, a new approach for protein representation learning based on the Siamese neural networks. We evaluate TripletProt comprehensively in protein functional annotation tasks including sub-cellular localization (14 categories) and gene ontology prediction (more than 2000 classes), which are both challenging multi-class multi-label classification machine learning problems. We compare the performance of TripletProt with the state-of-the-art approaches including recurrent language model-based approach (i.e., UniRep), as well as protein-protein interaction (PPI) network and sequence-based method (i.e., DeepGO). Our TripletProt showed an overall improvement of F1 score in the above mentioned comprehensive functional annotation tasks, solely relying on the PPI network. TripletProt and in general Siamese Network offer great potentials for the protein informatics tasks and can be widely applied to similar tasks.

## 1. Introduction

Proteins are the most important macromolecules in living systems with vital functions in virtually all biological systems and processes (Berg, Tymoczko, and Stryer 2012). Protein engineering has great potential to revolutionize medicine in the near future (Alley et al. 2019). Rational engineering approaches, in contrast to traditional methods, produce quantitative models of protein features. Although numerous properties and characteristics can be considered for protein engineering, protein functions mainly originate from a set of shared fundamental features (Alley et al. 2019). Current computational approaches aim to quantitatively model or analyze a small set of these features. Structure-based methods for protein function prediction conduct statistical analysis and simulate molecular dynamics based on free energy and thermostability (Rohl et al. 2004;). The scarcity of structural data along with computational intractability can be considered among the most important challenges these methods are faced with. On the other hand, data-driven co-evolutionary approaches are only promising in specific domains and in the presence of evolutionary information (Riesselman, Ingraham, and Marks 2018). Thus scalable methods are needed to address more important protein features, where a holistic representation can facilitate a domain-independent protein feature extraction.

Deep learning approaches achieving state-of-the-art performance in different machine learning applications are able to uncover representations in an end-to-end manner replacing classical feature engineering. Protein structure prediction (AlQuraishi 2019), (Asgari et al. 2019), function prediction (X. Liu 2017; Asgari and Mofrad 2015; Asgari, McHardy, and Mofrad 2019; Zhou et al. 2019) and semantic search (Schwartz et al. 2018) are promising state-of-the-art applications of deep networks in protein informatics. All of these approaches are, nonetheless, domain-specific and few studies have proposed a universal representation for proteins (Alley et al. 2019; Asgari and Mofrad 2015) as a challenging task. Proposing a comprehensive representation of proteins can significantly improve our understanding and prediction of the characteristics and functions of proteins.

Recently, pretrained representations have gained attention in various machine learning applications, especially for the language data. Vector representation of words in natural language processing (NLP) is a valuable approach to represent semantic and syntactic aspects of a word helping down-stream natural language understanding tasks at the low expense of raw text data (Collobert et al. 2011). Word embedding trained on language modeling of raw sequences/sentences projecting words to n-dimensional vectors, where similar words have close vectors (Mikolov et al. 2013). State-of-the-art approaches, BERT (Devlin et al. 2019) and XLNet (Yang et al. 2020), present a bidirectional language modeling which is inspired by denoising autoencoders. These methods involve considerable computational costs for training the model, hence motivating alternative approaches for representation learning.

Language modeling approaches on proteins (Rao et al. 2019;Heinzinger et al. 2019) often utilize protein sequences as an analogy to sentences in NLP. In the present work, we adopt a new perspective by exploiting the associations between proteins.

To capture the connectivity patterns observed in networks, node2vec (Grover and Leskovec 2016) utilizes a new notion of the neighborhood. In this method, nodes in the graph of proteins are mapped onto a low-dimensional space of features preserving relative distances among neighbors in the embedding graph. Node2vec uses a random-walk procedure to explore node neighbors. In the present study, we propose a novel embedding method for the representation of proteins based on protein-protein interaction networks. We focus on learning a new representation for proteins based on the Siamese neural networks (Bromley et al. 1993; Chopra, Hadsell, and LeCun 2005; Taigman et al. 2014). Specifically, we utilize a functional protein association network that integrates various sources of evidence concerning protein-protein interactions. Evidently, functional interactions of proteins are the backbone of the cellular machinery. Utilizing these interactions would be, therefore, invaluable to drawing new representations for proteins and understanding their roles in the mechanisms and functions of how cells work. In this study, we do not use the protein sequences in the creation of protein representations. Surprisingly, the new representations, trained only based on functional protein association networks, outperform the state-of-the-art approaches for the evaluated applications. Integration of protein sequences in the embedding process will likely achieve more precise representations, which will be the subject of future works.

## 2. Materials and Methods

### 2.1. Datasets

#### 2.1.1. STRING database

The STRING database (Szklarczyk et al. 2019) integrates and scores all publicly available sources of protein-protein associations including direct physical interaction, as well as indirect functional associations. In general, certain associations may indicate that proteins jointly contribute to a shared function but this does not necessarily imply they have direct binding. To achieve a comprehensive network, STRING complement this information with computational predictions. Various types of associations used by STRING to generate the interaction network are summarized in Figure 1.

**Figure 1.**
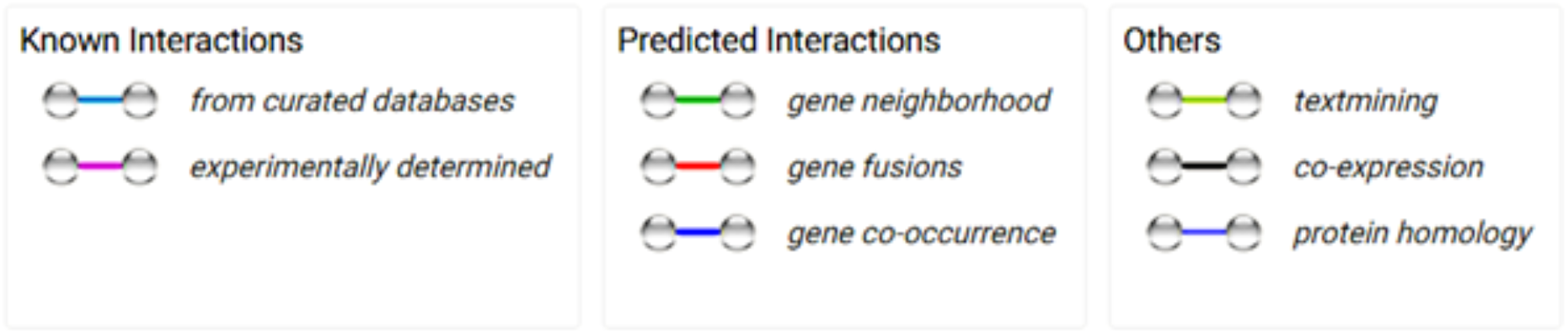
A summary of various types of evidence used for generating the protein-protein association network (Generated by STRING)

The most recent version of STRING (11.0) (Szklarczyk et al. 2019) covers proteins of 5090 distinct organisms. In the present study, we focus on 19,354 human proteins.

STRING annotates each protein-protein interaction with one or more ‘scores’, which are defined as indicators of confidence not necessarily the strength of the interaction. All scores range from 0 to 1, where 1 is the highest possible confidence. The 19,354 human proteins used in this study participate in more than 12 million interactions with various confidence scores. The interactions histogram of different confidence scores is presented in Figure 2, where all the associations are included. In the conducted experiences, we noticed that defining a threshold for the score leads to a poor classification performance. However, in the presence of a hardware bottleneck, a score threshold can be considered at the cost of performance loss.

**Figure 2.**
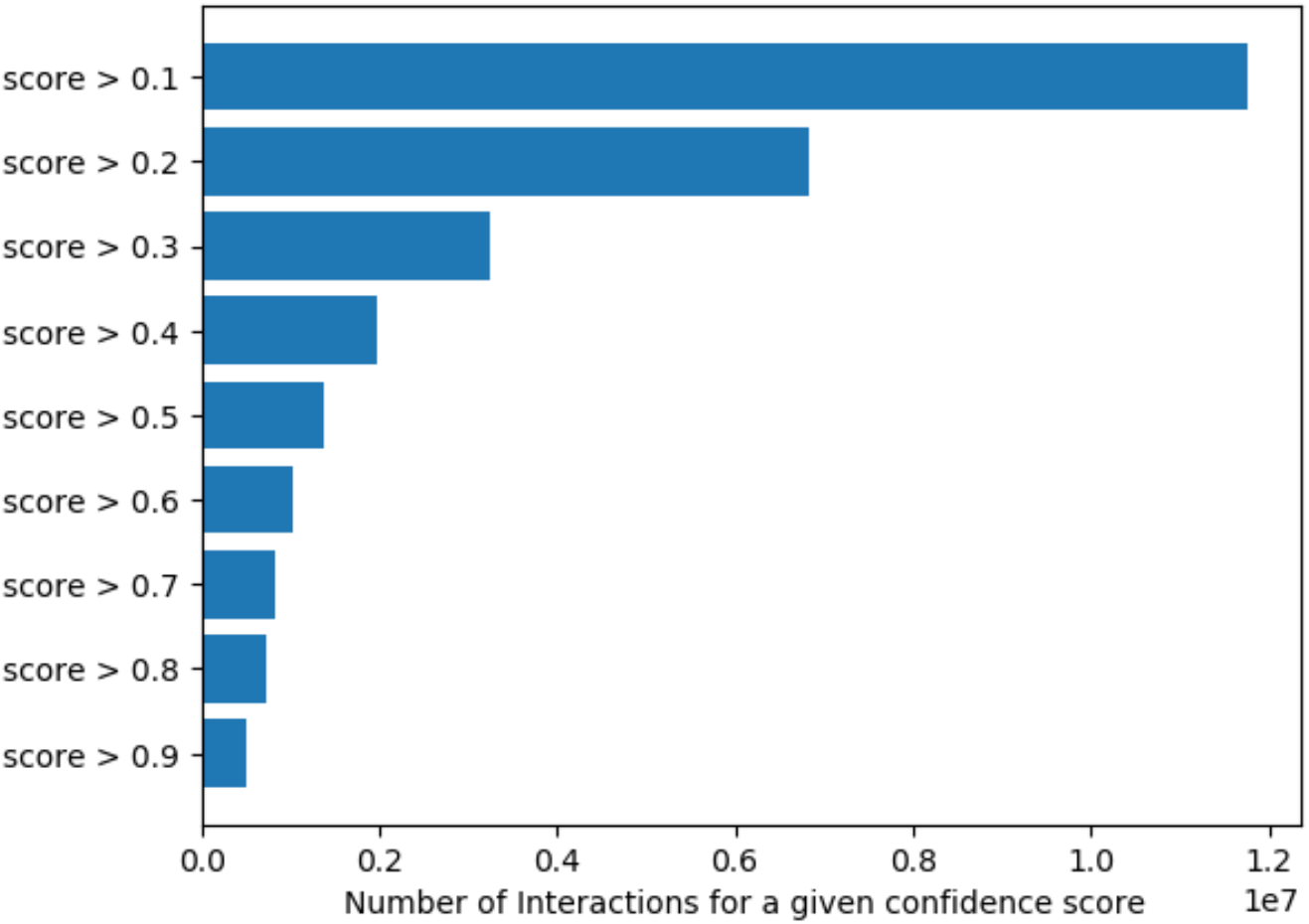
Number of interactions by changing the threshold score [0.1, 0.9]. STRING annotates each interaction with scores, i.e. how likely STRING judges an interaction to be true, as indicators of confidence not necessarily the strength of the interaction. All scores range from 0 to 1, where 1 is the highest possible confidence.

### 2.2. Proposed Method

We present an embedding approach to encode every protein in the association network into an n-dimensional vector. Generated embeddings as 64-dimensional feature vectors can be general representations of proteins. To evaluate the performance of these embeddings, they are employed as feature vectors in two examples, namely: (i) classifying protein subcellular localization and (ii) protein function prediction without the use of any other feature. The workflow of the proposed method is depicted in Figure 3.

**Figure 3.**
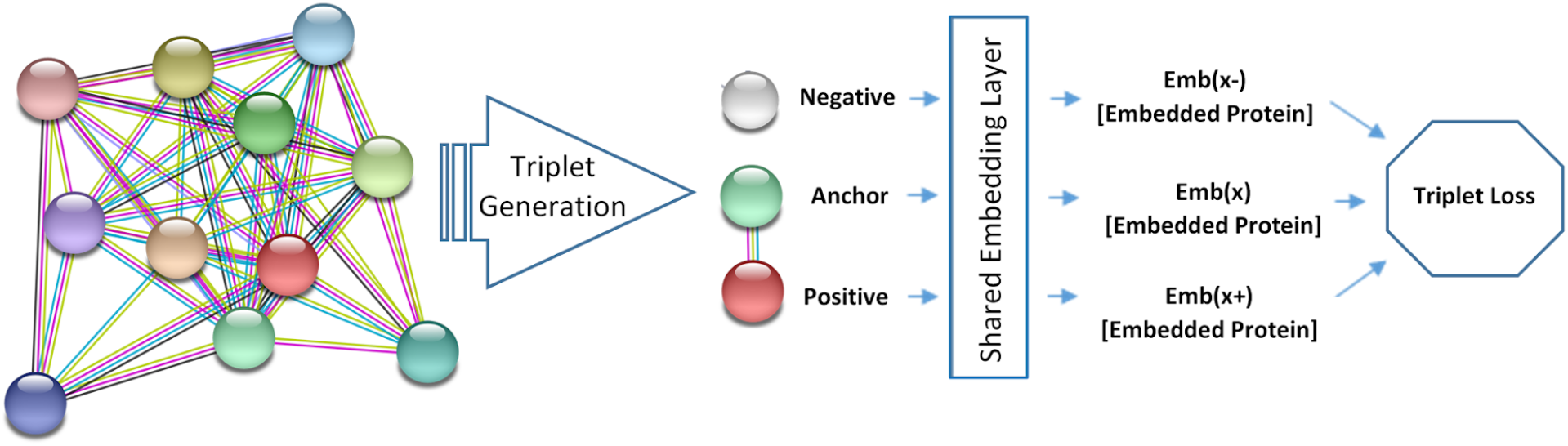
The proposed architecture to generate protein embeddings.The network of PPIs extracted from STRING was used to generate triplets. Each interaction pair is used to form a triplet. Each triplet is formed by x,x+,x−, where interacting proteins are considered as an anchor (x) and positive (x+). The embedding layer has shared parameters and generates representations denoted by Emb(x).(Sample network is generated by the STRING)

We train a variant of the Siamese Network, in which a triplet loss over the generated representation is adopted. Triplet networks (Hoffer and Ailon 2018) are inspired by Siamese (twin) networks. The embedding layer in the network receives triplets formed by anchor, positive and negative proteins (pr, pr+, and pr−). The network of protein-protein interactions (PPIs) extracted from STRING was used to generate triplets. Each interaction pair in the PPI network is used to form a triplet. Each triplet is formed by pr, pr+, pr−, where interacting proteins are considered as an anchor (pr), positive (pr+) and negative. Here, a random protein is considered as negative. The negative sample should interact with none of the two other interacting proteins.

The embedding layer has shared parameters and generates representations denoted by Emb(pr). This layer gets the indices of anchor, positive, and negative proteins as a triplet and generates 64-dimensional representations for these proteins. The triplet loss receives the embedded representations and computes the loss based on Equation (1) as follows:

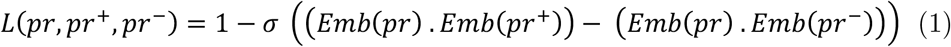

Where *σ* is the sigmoid function. As constructed, the triplet loss favors a small distance between the anchor and positive proteins, and a large distance for a pair of anchor and negative proteins. The shared embedding layer is used for generating embedding vectors for proteins. Since a single embedding is desired for the proteins, shared weights for the anchor, positive and negative proteins are enforced. The output size of the embedding layer is equal to 50. In other words, this is the size of the latent feature vector of each protein, which is the smallest and the most efficient embedding vector size for proteins in comparison to the state-of-the-art protein representations (Alley et al. 2019; Kulmanov, Khan, and Hoehndorf 2018). Current protein embedding techniques require far bigger vectors to reach acceptable performance.

To evaluate generated embeddings, a simple feedforward network receiving the embedding as input and predicting the target label is used. The utilized network architecture as the classifier can be significantly improved. However, to highlight the quality of the generated representation, we make use of a simple classifier commonly used in NLP for the evaluation of representations (Asgari and Mofrad 2015; Kiros et al. 2015).

## 3. Experiments

To evaluate the performance of our proposed deep representation learning approach, nicknamed TripletProt, we use it to produce representations in two different examples, namely: (i) the prediction of subcellular location; and (ii) protein function prediction, and compare the performance of our proposed method with the state-of-the-art approaches. Specifically, we evaluate the trained embedding in this study with those produced by two recent approaches: (1) DeepGO (Kulmanov, Khan, and Hoehndorf 2018) using sequence and network information (PPI network from the STRING database); and (2) UniRep (Alley et al. 2019), as a second baseline representation. DeepGo is a state-of-the-art method using a deep ontology classifier for prediction of protein functions from sequence and protein-protein interactions. UniRep is a state-of-the-art unified representation for proteins that can be applied for protein engineering informatics.

UniRep, generates embeddings in three different vector sizes (64, 256, and 1900). In this study, we have selected the largest Unirep vectors (1900). Our proposed approach outperformed UniRep while using a ~30 times smaller vector size (64). Furthermore, we have used only PPIs to train these embeddings taking less than 5 minutes with a basic computational setup.

### 3.1. Predicting subcellular location

The important role of subcellular location of proteins in basic science of cell functional machinery and applied targets such as drug design and protein functional annotation motivates the development of accurate machine learning methods for protein subcellular location prediction. Subcellular location prediction is currently an active area of research in protein informatics (Shen, Tang, and Guo 2019; Tung et al. 2017).

#### 3.1.1. Comparison with Multi-Kernel SVM

Shen (Shen, Tang, and Guo 2019) proposed a multi-kernel SVM to predict both multi-location and single-location proteins. They employed a position-specific scoring matrix (PSSM) and incorporated the physicochemical properties of proteins as features.

Table 1 summarizes the dataset used in this study collected from the Swiss-Prot database (release 55.3). This dataset contains 3106 samples, divided into 14 classes where 2580 proteins belong to one subcellular location, 480 to two locations, 43 to three locations, and 3 to four locations (Shen, Tang, and Guo 2019). Locative protein refers to a location assigned to a protein, since more than one location can be assigned to one protein, the total number of actual proteins is less than the number of locative proteins.

**Table 1.** Statistics of the dataset used by Multi-Kernel SVM

| Label | Subcellular localization | Number of sequences |
| --- | --- | --- |
| 1 | Centriole | 77 |
| 2 | Cytoplasm | 817 |
| 3 | Cytoskeleton | 79 |
| 4 | Endoplasmic reticulum | 229 |
| 5 | Endosome | 24 |
| 6 | Extracell | 385 |
| 7 | Golgi apparatus | 161 |
| 8 | Lysosome | 77 |
| 9 | Microsome | 24 |
| 10 | Mitochondrion | 361 |
| 11 | Nucleus | 1021 |
| 12 | Peroxisome | 47 |
| 13 | Plasma membrane | 354 |
| 14 | Synapse | 22 |
| No. of locative proteins | 3681 |  |
| No. of actual proteins | 3106 |  |

To evaluate the performance of our proposed approach, a five-fold cross-validation is conducted according to the reference papers. Average precision of the method is measured based on (1):

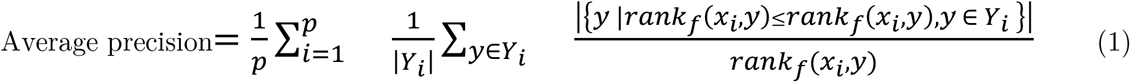

where p represents the number of samples, *rank*_*f*_(*x*, *y*) indicates the rank of y in Y on the descending order included from f (x, ·), and f (·, ·) represents the score returned by the classifier, and Y represents the real labels of sample x.

The features for the Multi-kernel SVM (see Table 2) are employed as described in (Shen, Tang, and Guo 2019): (i) the pseudo-position specific scoring matrix (PsePSSM) with 620-dimensions inspired by Chou’s pseudo amino acid (PseAAC) (Chou 2001) and obtained from the PSSM matrix; (ii) the pseudo-physicochemical properties (PsePP) of 1767 dimensions; (iii) the Average Blocks (AvBlock) introduced in (Jong Cheol Jeong, Xiaotong Lin, and Xue-Wen Chen 2011) is applied over PSSM and PsePP matrices to generate PSSM−AvBlock and PP−AvBlock features with 400 and 1140 dimensions respectively; (iv) PSSM−DWT is a 1040-dimensional feature for the PSSM matrix; and (v) PP–DWT is a 2964-dimensional feature for the PP matrix, where both are generated by applying the algorithm proposed in (Nanni, Lumini, and Brahnam 2014) based on Discrete Wavelet Transform (DWT) to extract the frequency and location information.

**Table 2.**
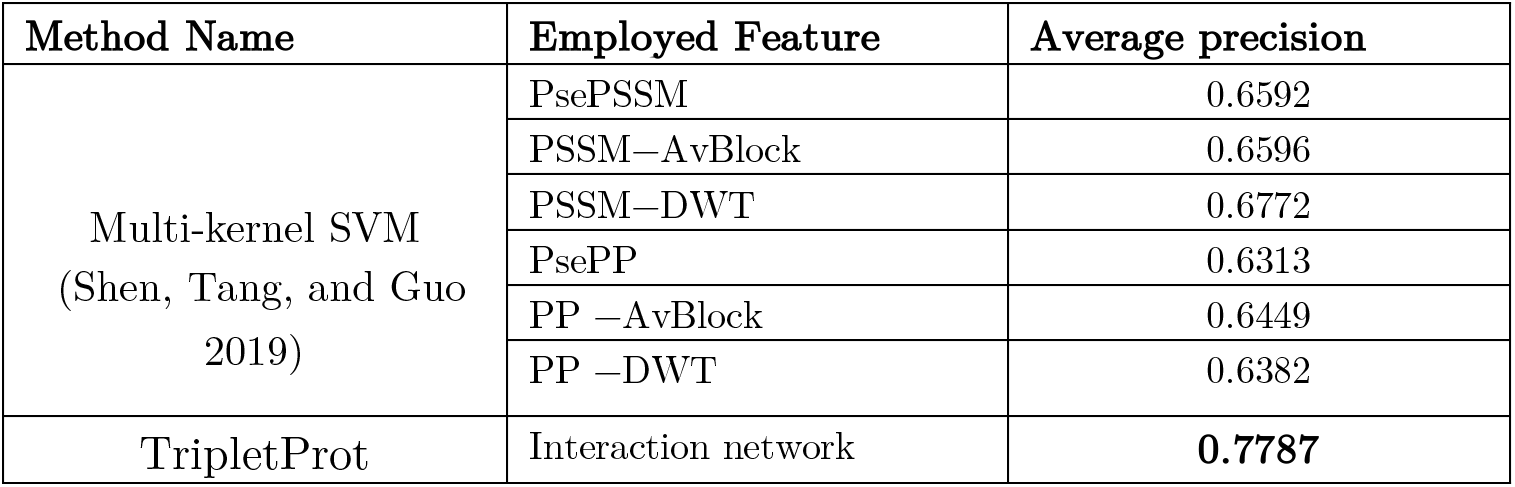
Performance of the proposed approach using five-fold cross validation

| Method Name | Employed Feature | Average precision |
| --- | --- | --- |
| Multi-kernel SVM<br>(Shen, Tang, and Guo<br>2019) | PsePSSM | 0.6592 |
|  | PSSM–AvBlock | 0.6596 |
|  | PSSM–DWT | 0.6772 |
|  | PsePP | 0.6313 |
|  | PP –AvBlock | 0.6449 |
|  | PP –DWT | 0.6382 |
| TripletProt | Interaction network | **0.7787** |

We show that our proposed approach without using any sequence feature significantly outperforms the state-of-the-art methods. Thanks to the precise representation of proteins trained over the protein-protein association network, no further extra sequence feature is required.

#### 3.1.2. Comparison with REALoc

REALoc (Tung et al. 2017) predicts subcellular localization of singleplex and multiplex proteins based on heterogeneous frameworks integrating machine learning methods using sequence-based features (e.g., amino acid composition, surface accessibility, and weighted sign aa index). REALoc consists of two layers. The first layer uses 32 SVM models and the second layer uses SVM results as input data to predict the six subcellular locations using a genetic algorithm optimized neural network (GANN) and majority voting (Vote). The results and statistics of the utilized dataset are summarized in Table 3.

**Table 3.** The number of proteins in the different subcellular locations used by REALoc

| Subcellular localization | Training dataset | Testing dataset |
| --- | --- | --- |
| Cell membrane | 1453 | 221 |
| Cytoplasm | 1542 | 197 |
| ER/Golgi | 562 | 136 |
| Mitochondrion | 462 | 133 |
| Nucleus | 2064 | 152 |
| Extracellular | 795 | 82 |
| Total | 6878<sup>a</sup> (5939<sup>b</sup>) | 925<sup>a</sup> (868<sup>b</sup>) |
| <sup>a</sup>Total number of proteins in all locations |  |  |
| <sup>b</sup>Total number of different proteins |  |  |

Each protein can be located in more than one subcellular location (see Figure 4), where for the sake of a clearer visualization the extracellular location is not shown. Similar to the previous experiment, a multi-label classification approach needs to be applied on this dataset, as every protein may potentially have multiple labels. Note that the labels here are designating different cellular compartments. For instance, in Figure 4 there are 34 proteins that are at the same time assigned to the “Cell membrane”, “Nucleus”, and “Cytoplasm”.

**Figure 4.**
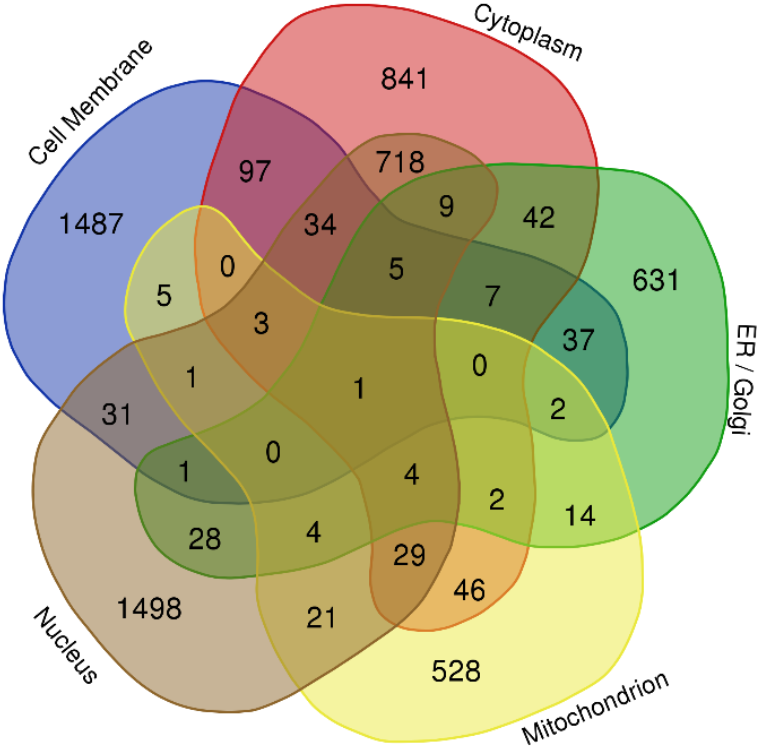
The Venn diagram showing the subcellular location overlaps for the proteins used by REALoc.

To evaluate our proposed methods, we conduct a five-fold cross-validation and, similar to the original study, we use the absolute true success rate, ATSR, based on the following equation (2).

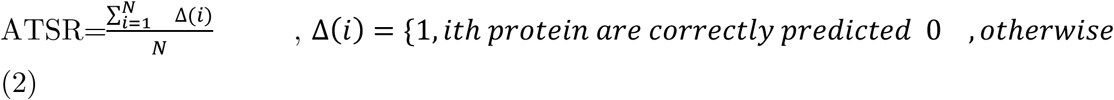

ATSR is an evaluation metric requiring correct prediction for all subcellular locations of a protein. Of course, ATSR is a less forgiving metric in comparison to other metrics. ATST considers a prediction correct if and only if all of its subcellular positions are correctly predicted. Evaluation results for train dataset using the five-fold cross-validation are reported in Table 4 and on the test set in Table 5, respectively.

**Table 4.** The performance of the proposed approach in comparison with REALoc using five-fold cross-validation and ATST score for the 5939p dataset is presented in the table.

| Subcellular localization | REALoc_Vote | REALoc_GANN | Proposed Approach |
| --- | --- | --- | --- |
| Cell membrane | 79.6 | 80.6 | 80.7 |
| Cytoplasm | 63.5 | 65.6 | 75.0 |
| ER/Golgi | 74.4 | 81.1 | 64.9 |
| Mitochondrion | 76.0 | 75.3 | 80.3 |
| Nucleus | 72.7 | 74.8 | 86.4 |
| Extracellular | 76.5 | 80.9 | 74.5 |
| Overall | 72.8 | 75.3 | **77.0** |

**Table 5.** The performance of the proposed approach in comparison with REALoc for the 868p testing dataset.

| Subcellular localization | REALoc_Vote | REALoc_GANN | TripletProt |
| --- | --- | --- | --- |
| Cell membrane | 57.9 | 47.5 | 81.0 |
| Cytoplasm | 57.4 | 50.8 | 82.0 |
| ER/Golgi | 53.7 | 52.9 | 51.5 |
| Mitochondrion | 53.4 | 54.1 | 53.5 |
| Nucleus | 58.3 | 55.8 | 71.8 |
| Extracellular | 80.0 | 80.0 | 57.1 |
| Overall | 57.1 | 52.5 | **66.1** |

As presented in Table 4 and 5, although the results of the five-fold cross-validation are not consistently improved on both cross-validation and test set, the overall results are considerably improved in both cases.

#### 3.1.3. Visual comparison with UniRep using t-SNE

In this section, a qualitative comparison of the tripletProt with the uniRep is performed based on the 2D representations generated by t-SNE (Figure 5).

**Figure 5.**
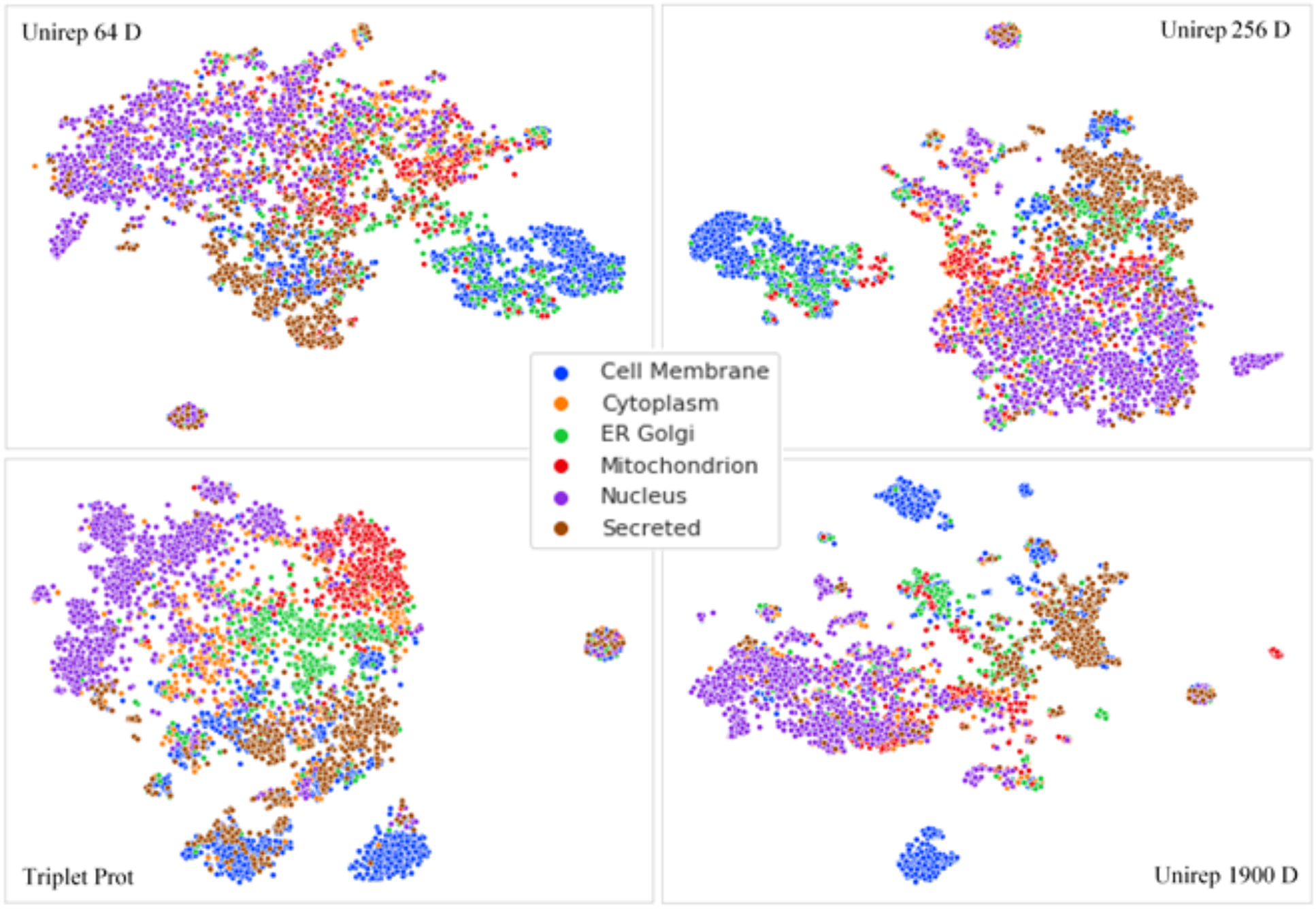
t-SNE visualizations produced using various dimensions of the UniRep in comparison with the Triplet Prot

Among three different vector dimensions of the uniRep, the best separation is generated using the most elaborate one. However, as shown in Figure 5, the separability of the representation trained using 1900D vectors is lower than the TripletProt. The subcellular components are considerably more disjointed for the vectors trained using TripletProt compared to the representations trained by uniRep.

### 3.2. Computational function prediction of proteins

As the last set of experiments, we utilize the trained embeddings in machine learning for protein function annotation. Experimental methods for protein function prediction are time-consuming and expensive and have therefore been conducted only for a limited set of proteins. Gene Ontologies (GO) characterize protein functions using more than 40,000 classes categorized into three sub-ontologies: (i) Molecular Function (MF), (ii) Biological Process (BP), and (iii) Cellular Component (CC). Protein function prediction is a multi-class, multi-label problem, as each protein may have multiple distinct functions. Details of the employed dataset are summarized in Table 6. DeepGo splits proteins into a training set (80%) and testing set (20%). GO classes are ranked based on their number of annotations and the top 932 terms for BP, 589 terms for MF and 436 terms for the CC ontology are selected.

**Table 6.** Dataset used for protein function prediction

| Sub Ontology | Number of Terms (Classes) | Minimum Number of Proteins annotated by each Term |
| --- | --- | --- |
| BP - Biological Processes | 932 | 250 |
| MF - Molecular Functions | 589 | 50 |
| CC - Cellular Components | 436 | 50 |

To conduct a fair comparison, we adopt the same train and test datasets as used in the DeepGo study. The label vector of each protein is prepared based on the annotated GO terms. Therefore, the length of the label vector is equal to the number of GO terms. A binary vector with the entries of 1 if the protein is annotated with the corresponding term and 0 otherwise. Every sub-ontology has distinct label vectors that are evaluated in three sets of experiments.

DeepGO creates a multi-species knowledge graph based on protein-protein interactions. Where two types of edges are considered for inter-species orthologous proteins or intra-species interactions. After generating graph embedding, the final representation vector for each protein is produced by integrating the sequence representation. As a result, DeepGO produces a multi-modal representation that employs both protein interactions and their sequences. For the comparison in this study, we use the same embedding vectors shared by DeepGO.

UniRep generates protein representations using a recurrent neural network and about 24 million primary amino acid sequences. Their model is trained to predict the next amino acid minimizing the cross-entropy loss and consequently learn how to represent proteins. They trained 1900-dimension embeddings for about 3 weeks on 4 Nvidia K80 GPUs. Two variants of the embeddings are also available with sizes 256 and 64. In this study, we utilize the 1900-dimension embeddings achieving the best results in the original UniRep paper.

Table 7 presents the average precision results obtained by the embeddings trained over the proposed approach as compared with two state-of-the-art embedding approaches, namely DeepGO and UniRep.

**Table 7.** Average precision results produced by DeepGO vectors, Unirep representations and our proposed embeddings. The same neural network is used as the classifier for all three compared representations. Statistics of proteins and GO terms are described in Table 6.

| GO-Category | Deep-GO | Unirep | TripletProt |
| --- | --- | --- | --- |
| BP - Biological Processes | 0.471 | 0.428 | **0.508** |
| MF - Molecular Functions | 0.517 | 0.522 | **0.530** |
| CC - Cellular Components | 0.677 | 0.678 | **0.703** |

Table 7 shows the results of comparing three recent protein embedding approaches with our method using the same classifier for all. A vanilla convolutional network is utilized in this study as the classifier to exhibit the strength of the proposed embeddings.

A significant improvement is achieved here for predicting BP terms, particularly as compared to the prediction by UniRep. UniRep, which does not make any use of protein interaction information, tends to exhibit poorly in comparison to our proposed method and DeepGo, highlighting the critical importance of the PPI information in functional prediction of proteins.

In the next set of experiments, another well-known metric is adopted for performance evaluation. F1 takes both precision and recall into account by taking their harmonic average.

Since protein function prediction is a multilabel, multiclass classification problem, the weights of different classes in the final metric computation can be determined using two variations of F1. The first mode, called macro averaging of the F1, assigns equal weight to all classes, regardless of having an imbalanced data set. On the other hand, micro averaging assigns larger weights to the classes having more samples. The micro F1, macro F1, and average precision results of state-of-the-art approaches along with our TripletProt are presented in Figures 5-7, respectively. It is important to note that F1 needs a threshold to be computed in contrast with average precision, which considers the rank of predicted samples and is not dependent on a specific threshold value. To evaluate the effect of a threshold, experiments are conducted for different threshold values in the range of 0.05 and 0.5.

As shown in Figures 6-8, the best results are achieved by the TripletProt for all categories of the GO terms. UniRep and DeepGo achieved similar results for molecular function and cellular compartment terms. DeepGo is the second best after TripletProt for the biological process terms, likely because of incorporating the PPI network in contrast to UniRep, which solely relies on the sequence information.

**Figure 6.**
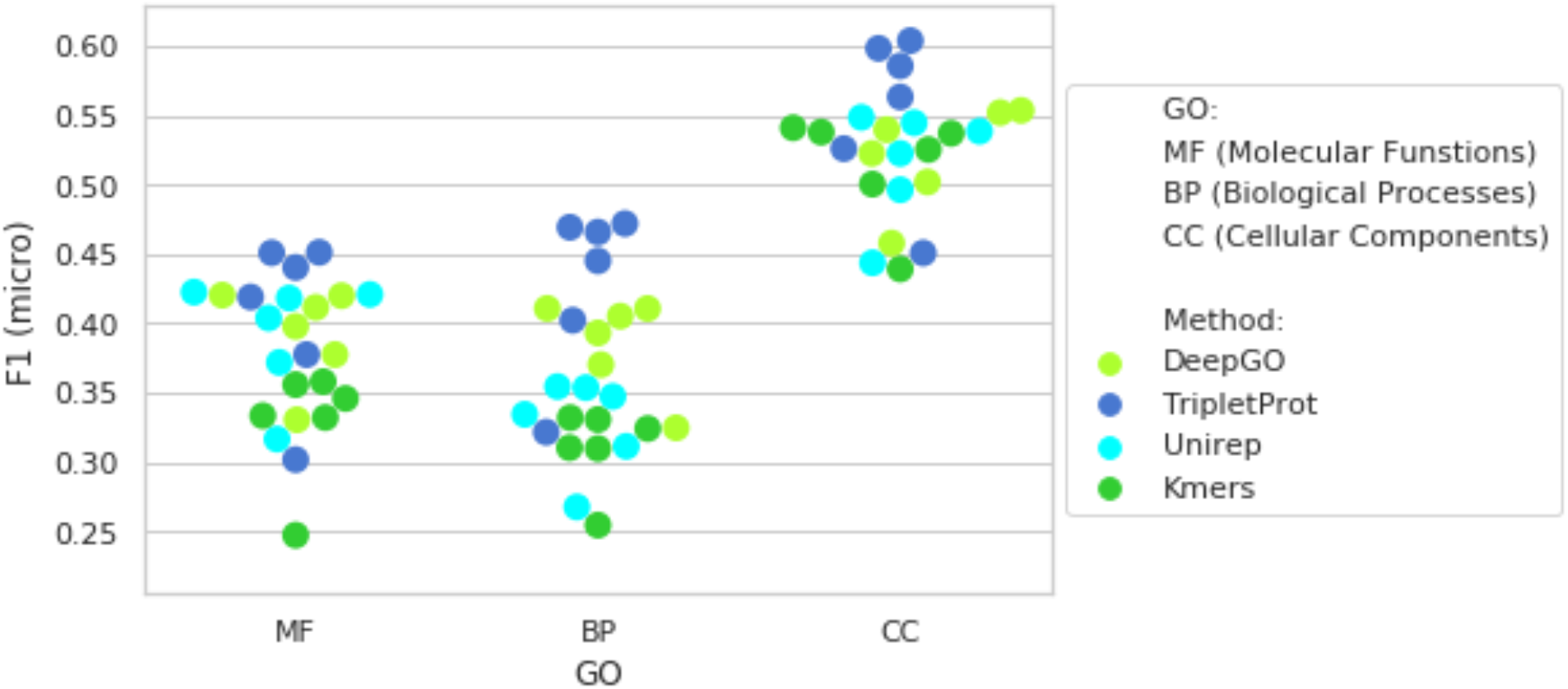
F1 (micro) results of the three compared method over the different categories of the GO terms repeated using 10 various thresholds in range (0.05, 0.5, step:0.05)

For each of the three GO categories, there are hundreds of different sub-categories. Macro averaging for the F1 metric, assigns equal weights to all terms. Distribution of the samples is heavily imbalanced for different terms, hence enforcing equal weights for all the terms is not likely justified for this experiment. It is common, however, to have no predicted sample for specific classes, resulting in zero F1 and significantly affecting the final average. TripletProt outperforms the state-of-the-art approaches in Gene ontology prediction tasks with an exception of MF categories. Even in MF predictions, the results are comparable to the other approaches.

The most stable results are achieved based on the average precision metric (see Figure 8). Exact predicted scores are used only for ranking the results. Therefore, specific thresholds are not required as a borderline between positive and negative decisions. As shown in Figure 7, the results are not dependent on various threshold values. TripletProt outperforms the other approaches even for the MF terms.

**Figure 7.**
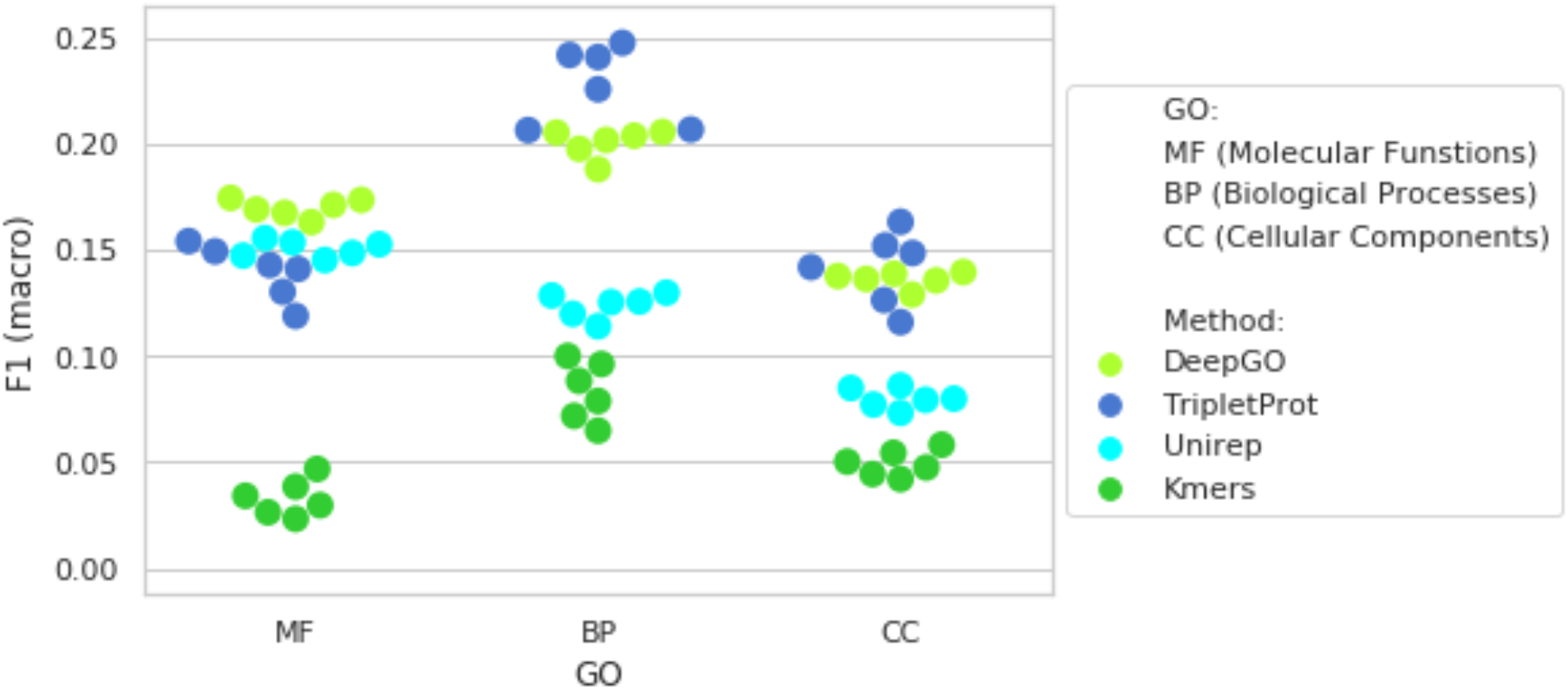
F1 (macro) results of the three compared method over the different categories of the GO terms repeated using 10 various thresholds in range (0.05, 0.5, step: 0.05)

**Figure 8.**
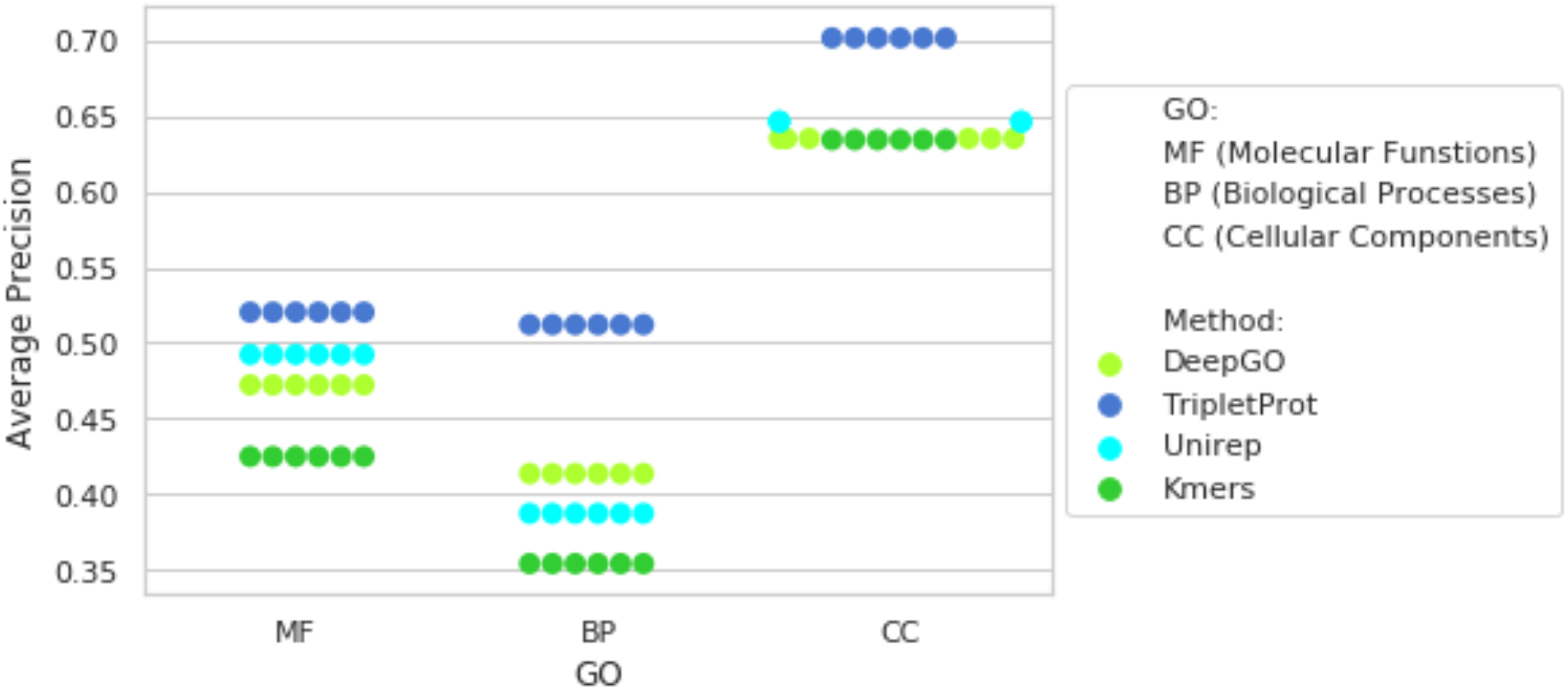
The average precision results of the three compared methods over the different categories of the GO terms repeated using 10 various thresholds in range (0.05, 0.5, step:0.05) are shown

## Conclusion

In this paper, we presented TripletProt, a novel Siamese neural network-based approach for protein representation learning. The most important distinction of our proposed method is relying on the PPI network. State-of-the-art approaches typically consider only protein sequences or a combination of sequence and PPI network. Significance of the achieved improvements seems to be notable especially for proteins without known sequences. The computational cost of the generated representations for any potential application is significantly lower than comparable methods since the length of the representations is significantly smaller than that in other approaches. Two sample applications were used to evaluate the representations generated by the TripletProt. Specifically, subcellular protein location and function prediction of proteins as multi-class multi-label classification problems were formulated and evaluated in comparison with state-of-the-art methods. Significant improvements were achieved in most of the conducted experiments. In future work, we will integrate protein sequences in the embedding process to achieve more precise representations.

## References

Alley, Ethan C., Grigory Khimulya, Surojit Biswas, Mohammed AlQuraishi, and George M. Church. 2019. “Unified Rational Protein Engineering with Sequence-Based Deep Representation Learning.” Nature Methods 16 (12): 1315–22. https://doi.org/10.1038/s41592-019-0598-1.

AlQuraishi, Mohammed. 2019. “End-to-End Differentiable Learning of Protein Structure.” Cell Systems 8 (4): 292–301.e3. https://doi.org/10.1016/j.cels.2019.03.006.

Asgari, Ehsaneddin, Alice C McHardy, and Mohammad RK Mofrad. 2019. “Probabilistic Variable-Length Segmentation of Protein Sequences for Discriminative Motif Discovery (DiMotif) and Sequence Embedding (ProtVecX).” Scientific Reports 9 (1): 3577.

Asgari, Ehsaneddin, and Mohammad R. K. Mofrad. 2015. “Continuous Distributed Representation of Biological Sequences for Deep Proteomics and Genomics.” Edited by Firas H Kobeissy. PLOS ONE 10 (11): e0141287. https://doi.org/10.1371/journal.pone.0141287.

Asgari, Ehsaneddin, Nina Poerner, Alice C. McHardy, and Mohammad R. K. Mofrad. 2019. “DeepPrime2Sec: Deep Learning for Protein Secondary Structure Prediction from the Primary Sequences.” BioRxiv, July, 705426. https://doi.org/10.1101/705426.

Berg, Jeremy M., John L. Tymoczko, and Lubert Stryer. 2012. Biochemistry. 7th ed. New York: W.H. Freeman.

Bromley, Jane, James W. Bentz, Léon Bottou, Isabelle Guyon, Yann Lecun, Cliff Moore, Eduard Säckinger, and Roopak Shah. 1993. “SIGNATURE VERIFICATION USING A ‘SIAMESE’ TIME DELAY NEURAL NETWORK.” International Journal of Pattern Recognition and Artificial Intelligence 07 (04): 669–88. https://doi.org/10.1142/S0218001493000339.

Chopra, S., R. Hadsell, and Y. LeCun. 2005. “Learning a Similarity Metric Discriminatively, with Application to Face Verification.” In 2005 IEEE Computer Society Conference on Computer Vision and Pattern Recognition (CVPR’05), 1:539–46. San Diego, CA, USA: IEEE. https://doi.org/10.1109/CVPR.2005.202.

Chou, Kuo-Chen. 2001. “Prediction of Protein Cellular Attributes Using Pseudo-Amino Acid Composition.” Proteins: Structure, Function, and Genetics 43 (3): 246–55. https://doi.org/10.1002/prot.1035.

Collobert, Ronan, Jason Weston, Léon Bottou, Michael Karlen, Koray Kavukcuoglu, and Pavel Kuksa. 2011. “Natural Language Processing (Almost) from Scratch.” Journal of Machine Learning Research 12 (Aug): 2493–2537.

Devlin, Jacob, Ming-Wei Chang, Kenton Lee, and Kristina Toutanova. 2019. “BERT: Pre-Training of Deep Bidirectional Transformers for Language Understanding.” ArXiv:1810.04805 [Cs], May. http://arxiv.org/abs/1810.04805.

Grover, Aditya, and Jure Leskovec. 2016. “Node2vec: Scalable Feature Learning for Networks.” ArXiv:1607.00653 [Cs, Stat], July. http://arxiv.org/abs/1607.00653.

Heinzinger, Michael, Ahmed Elnaggar, Yu Wang, Christian Dallago, Dmitrii Nechaev, Florian Matthes, and Burkhard Rost. 2019. “Modeling Aspects of the Language of Life through Transfer-Learning Protein Sequences.” BMC Bioinformatics 20 (1): 723. https://doi.org/10.1186/s12859-019-3220-8.

Hoffer, Elad, and Nir Ailon. 2018. “Deep Metric Learning Using Triplet Network.” ArXiv:1412.6622 [Cs, Stat], December. http://arxiv.org/abs/1412.6622.

Jong Cheol Jeong, Xiaotong Lin, and Xue-Wen Chen. 2011. “On Position-Specific Scoring Matrix for Protein Function Prediction.” IEEE/ACM Transactions on Computational Biology and Bioinformatics 8 (2): 308–15. https://doi.org/10.1109/TCBB.2010.93.

Kiros, Ryan, Yukun Zhu, Ruslan Salakhutdinov, Richard S. Zemel, Antonio Torralba, Raquel Urtasun, and Sanja Fidler. 2015. “Skip-Thought Vectors.” ArXiv:1506.06726 [Cs], June. http://arxiv.org/abs/1506.06726.

Kulmanov, Maxat, Mohammed Asif Khan, and Robert Hoehndorf. 2018. “DeepGO: Predicting Protein Functions from Sequence and Interactions Using a Deep Ontology-Aware Classifier.” Edited by Jonathan Wren. Bioinformatics 34 (4): 660–68. https://doi.org/10.1093/bioinformatics/btx624.

Liu, Xueliang. 2017. “Deep Recurrent Neural Network for Protein Function Prediction from Sequence.” ArXiv:1701.08318 [Cs, q-Bio, Stat], January. http://arxiv.org/abs/1701.08318.

Mikolov, Tomas, Ilya Sutskever, Kai Chen, Greg S Corrado, and Jeff Dean. 2013. “Distributed Representations of Words and Phrases and Their Compositionality.” In Advances in Neural Information Processing Systems 26, edited by C. J. C. Burges, L. Bottou, M. Welling, Z. Ghahramani, and K. Q. Weinberger, 3111–3119. Curran Associates, Inc. http://papers.nips.cc/paper/5021-distributed-representations-of-words-and-phrases-and-their-compositionality.pdf.

Nanni, Loris, Alessandra Lumini, and Sheryl Brahnam. 2014. “An Empirical Study of Different Approaches for Protein Classification.” The Scientific World Journal 2014: 1–17. https://doi.org/10.1155/2014/236717.

Rao, Roshan, Nicholas Bhattacharya, Neil Thomas, Yan Duan, Xi Chen, John Canny, Pieter Abbeel, and Yun S. Song. 2019. “Evaluating Protein Transfer Learning with TAPE.” ArXiv:1906.08230 [Cs, q-Bio, Stat], June. http://arxiv.org/abs/1906.08230.

Riesselman, Adam J., John B. Ingraham, and Debora S. Marks. 2018. “Deep Generative Models of Genetic Variation Capture the Effects of Mutations.” Nature Methods 15 (10): 816–22. https://doi.org/10.1038/s41592-018-0138-4.

Rohl, Carol A, Charlie E.M Strauss, Kira M.S Misura, and David Baker. 2004. “Protein Structure Prediction Using Rosetta.” Elsevier Logo Journals & Books Esmaeil Nourani Methods in Enzymology 383: 66–93. https://doi.org/10.1016/S0076-6879(04)83004-0.

Schwartz, Ariel S, Gregory J Hannum, Zach R Dwiel, Michael E Smoot, Ana R Grant, Jason M Knight, Scott A Becker, Jonathan R Eads, Matthew C LaFave, and Harini Eavani. 2018. “Deep Semantic Protein Representation for Annotation, Discovery, and Engineering.” BioRxiv, 365965.

Shen, Yinan, Jijun Tang, and Fei Guo. 2019. “Identification of Protein Subcellular Localization via Integrating Evolutionary and Physicochemical Information into Chou’s General PseAAC.” Journal of Theoretical Biology 462 (February): 230–39. https://doi.org/10.1016/j.jtbi.2018.11.012.

Szklarczyk, Damian, Annika L Gable, David Lyon, Alexander Junge, Stefan Wyder, Jaime Huerta-Cepas, Milan Simonovic, et al. 2019. “STRING V11: Protein–Protein Association Networks with Increased Coverage, Supporting Functional Discovery in Genome-Wide Experimental Datasets.” Nucleic Acids Research 47 (D1): D607–13. https://doi.org/10.1093/nar/gky1131.

Taigman, Yaniv, Ming Yang, Marc’Aurelio Ranzato, and Lior Wolf. 2014. “DeepFace: Closing the Gap to Human-Level Performance in Face Verification.” In 2014 IEEE Conference on Computer Vision and Pattern Recognition, 1701–8. Columbus, OH, USA: IEEE. https://doi.org/10.1109/CVPR.2014.220.

Tung, Chi-Hua, Chi-Wei Chen, Han-Hao Sun, and Yen-Wei Chu. 2017. “Predicting Human Protein Subcellular Localization by Heterogeneous and Comprehensive Approaches.” Edited by Bin Liu. PLOS ONE 12 (6): e0178832. https://doi.org/10.1371/journal.pone.0178832.

Yang, Zhilin, Zihang Dai, Yiming Yang, Jaime Carbonell, Ruslan Salakhutdinov, and Quoc V. Le. 2020. “XLNet: Generalized Autoregressive Pretraining for Language Understanding.” ArXiv:1906.08237 [Cs], January. http://arxiv.org/abs/1906.08237.

Zhou, Naihui, Yuxiang Jiang, Timothy R Bergquist, Alexandra J Lee, Balint Z Kacsoh, Alex W Crocker, Kimberley A Lewis, George Georghiou, Huy N Nguyen, and Md Nafiz Hamid. 2019. “The CAFA Challenge Reports Improved Protein Function Prediction and New Functional Annotations for Hundreds of Genes through Experimental Screens.” BioRxiv, 653105.

